# Synaptic transmission modulates while non-synaptic processes govern the transition from pre-ictal to seizure activity in vitro

**DOI:** 10.1101/280321

**Authors:** Marom Bikson, Ana Ruiz-Nuño, Dolores Miranda, Greg Kronberg, Premysl Jiruska, John E Fox, John G.R. Jefferys

**Author notes:** Corresponding author: Marom Bikson. Current addresses: PJ, Department of Developmental Epileptology, Institute of Physiology, The Czech Academy of Sciences, Prague, Czech Republic JGRJ, Department of Pharmacology, University of Oxford, Oxford, UK.

## Abstract

It is well established that non-synaptic mechanisms can generate electrographic seizures after blockade of synaptic function. We investigated the interaction of intact synaptic activity with non-synaptic mechanisms in the isolated CA1 region of rat hippocampal slices using the “elevated-K^+^” model of epilepsy. Elevated K^+^ ictal bursts share waveform features with other models of electrographic seizures, including non-synaptic models where chemical synaptic transmission is suppressed, such as the low-Ca^2+^model. These features include a prolonged (several seconds) negative field shift associated with neuronal depolarization and superimposed population spikes. When population spikes are disrupted for up to several seconds, intracellular recording demonstrated that the prolonged suppression of population spikes during ictal activity was due to depolarization block of neurons. Elevated-K^+^ ictal bursts were often preceded by a build-up of “pre-ictal” epileptiform discharges that were characterized as either “slow-transition” (localized and with a gradual increase in population spike amplitude, reminiscent non-synaptic neuronal aggregate formation, presumed mediated by extracellular K+ concentrations ([K+])_o_ accumulation), or “fast-transition” (with a sudden increase in population spike amplitude, presumed mediated by field effects). When ictal activity had a fast-transition it was preceded by fast-transition pre-ictal activity; otherwise population spikes developed gradually at ictal event onset. Addition of bicuculline, a GABAA receptor antagonist, suppressed population spike generation during ictal activity, reduced pre-ictal activity, and increased the frequency of ictal discharges. Nipecotic acid and NNC-711, both of which block GABA re-uptake, increased population spike amplitude during ictal bursts and promoted the generation of preictal activity. By contrast, addition of ionotropic glutamate-receptor antagonists (NBQX, D-APV) had no consistent effect on ictal burst waveform or frequency and did not fully suppress pre-ictal activity. Similarly, CGP 55848, a GABA_B_ receptor antagonist, has no significant effect on pre-ictal activity or burst frequency (although it did increase burst duration slightly). Our results are consistent with the hypothesis that non-synaptic mechanisms underpin the generation of ictal bursts in CA1 and that GABA_A_ synaptic mechanisms can shape event development by delaying event initiation and counteracting depolarization block.

## Introduction

Focal epileptic seizures are usually explained by dysfunction of synaptically driven neuronal networks (Bradford, 1995). However, the generation of electrographic seizures after synaptic transmission was blocked in the hippocampal slice with low extracellular Ca2+ ^concentrations ([Ca^2^+]^o^)^, ^showing that synaptic function is not strictly necessary (Jefferys and^ Haas, 1982; Yaari et al., 1983). Additional *in vivo* (Feng and Durand, 2003) and *in vitro* (Bikson et al., 2002) epilepsy models based on suppressing synaptic function were developed, while in some epilepsy models with normally intact synaptic transmission seizures persisted after blocking synaptic function (Pumain et al., 1985; Jensen and Yaari, 1988; Patrylo et al., 1994; Demir et al., 1999). The importance of synapses in seizure generation is evidenced by anticonvulsants targeting synaptic transmission (Bialer et al., 2002) and other animal models of epilepsy (Traynelis and Dingledine, 1988; Leschinger et al., 1993; Barbarosie et al., 2002; Bragin et al., 2009; Zhang et al., 2012). Yet on their own non-synaptic mechanisms can be sufficient to initiate, synchronize, propagate, and terminate epileptic activity (Jefferys and Haas, 1982; Haas and Jefferys, 1984; Pumain et al., 1985; Bikson et al., 1999; Bikson et al., 2003a; Lian et al., 2003; Park and Durand, 2006). In those conditions where non-synaptic mechanisms are sufficient for electrographic seizures but synaptic transmission is intact, what are the relative roles of synaptic and non-synaptic processes?

The mechanism of transition from asymptomatic inter-ictal activity to symptomatic ictal seizures is one of the central goals of epilepsy research. Synaptic mechanisms facilitate inter-ictal epileptiform activity (de Curtis and Avanzini, 2001; Huberfeld et al., 2011; Fujita et al., 2014),but inter-ictal activity may be poorly correlated with seizure discharge severity, frequency, or anatomical focus (Gotman and Marciani, 1985; Swartzwelder et al., 1987; Bragdon et al., 1992; de Curtis and Avanzini, 2001). Indeed, in some epilepsy models, interictal epileptiform activity can suppress electrographic seizures (Swartzwelder et al., 1987; Jensen and Yaari, 1988; Bragdon et al., 1992; Barbarosie and Avoli, 1997). Studies in human epilepsy and animal models have suggested a sub-class of inter-ictal events: “pre-ictal” discharges which are more correlated temporally and spatially with seizure initiation and present distinct dependencies on synaptic function (Bragin et al., 2009; Cymerblit-Sabba and Schiller, 2010; Jiruska et al., 2010; Huberfeld et al., 2011; Zhang et al., 2012; Fujita et al., 2014; Perucca et al., 2014).

Thus, fundamental questions remain about the interaction of synaptic and non-synaptic communication in epilepsy. To address these questions, we used the elevated-K+ model of epilepsy in the CA1 region of rat hippocampal slices (Traynelis and Dingledine, 1988; Jensen and Yaari, 1997) where inhibitory and excitatory synaptic function is normally intact (Poolos and Kocsis, 1990; Jensen et al., 1993), yet non-synaptic interactions have been shown to be sufficient to generate electro-graphic seizures after synaptic transmission is pharmacologically blocked (Jensen and Yaari, 1988). CA1 discharges are modulated by interictal discharges in CA3(Traynelis and Dingledine, 1988; Jensen and Yaari, 1997), therefore, we disconnected these regions to remove a confounding variable. We selectively modulated excitatory and inhibitory synaptic pathways to identify the roles of synaptic transmission in the generation of ictal bursts in CA1. We propose that under conditions that support non-synaptic seizures synaptic function plays a modulatory role based in the conversion of preictal events. Under this framework, apparently counterintuitive responses for GABAergic and glutamatergic antagonists as well as diverse results on ictogenesis in CA1can be reconciled.

## Materials and Methods

Transverse hippocampal slices (350-400 µm) were prepared from male Sprague-Dawley rats (180-225g; anesthetized with ketamine/medetomidine or ketamine/xylazine). The slices were submerged in a holding chamber filled with “normal” ACSF gassed with 95% O_2_ 5% CO_2_, and comprising (in mM): 125 NaCl, 26 NaHCO_3_, 3 KCl, 2 CaCl_2_, 1.0 MgCl_2_, 1.25 NaH_2_PO_4_, and 10 glucose. After>60 min slices were transferred to an interface recording chamber. Complete mechanical lesions were made, in the CA2 region, across the Schaffer collateral pathway to promote the initiation of ictal events in CA1 (Jensen and Yaari, 1988) and to prevent complication in results interpretation due to modulation of inter-ictal activity in CA3 by drugs.

Spontaneous activity was induced by perfusion of slices (>60 min) with “elevated-K+” ACSF consisting of (in mM): 125 NaCl, 26 NaHCO_3_, 7.5-8 KCl, 1.0 CaCl_2_, 1.0-1.2 MgCl_2_, 1.25 NaH_2_PO_4_, and 10 glucose. These concentrations are comparable to the ionic constitution of the extracellular microenvironment during seizures *in vivo* (Somjen and Giacchino, 1985; Lux et al., 1986) and of the superfusates used in previous *in vitro* studies examining the effects of elevated K+ on prolonged electrographic seizures (Leschinger et al., 1993; Jensen and Yaari, 1997). Only slices generating “tonic ictal” activity, defined here as a continuous (though not necessarily flat) >2 mV negative shift in the field potential lasting > 5 seconds, were accepted in this study.

Conventional recording techniques were used to measure activity from the CA1 pyramidal cell region. Extracellular field potentials were recorded with glass micropipettes (2-8 MΩ) filled with ACSF. Intracellular glass micropipettes were filled with 2 M potassium methylsulphate and had resistances of 60-120 MΩ Neurons with <-55 mV (average -62 ± 5 mV) resting membrane potentials and action potentials > 60 mV were accepted in this study (Jensen and Yaari, 1997).

(±)-Nipecotic acid and (-)-Bicuculline methiodide were obtained from Sigma (Poole, U.K.). CGP 55845, NNC-711, NBQX and D-AP5 were obtained from Tocris (Bristol, UK). All drugs were applied via the superfusate for >20 min.

All signals were amplified and low-pass filtered (<3 kHz field, <10 kHz intracellular) with an Axoclamp-2B or 2A (Axon Instruments, Union City, U.S.A) and Neurolog NL-106 and NL-125 (Digitimer, Hertfordshire, U.K) or Cygnus FLA-01 (Cygnus Technology, Delaware Water Gap, PA) post-amplifiers. Signals were digitised using a Power 1401 and Signal/Spike software (Cambridge Electronic Design, Cambridge, U.K.). Results are reported as: mean ± standard deviation, n = number of slices or cells, as appropriate; changes were considered significant if p<0.05 using Student’s paired t-test. All procedures were approved by the University of Birmingham and the City College of New York animal welfare committees.

## Results

### General characteristics of elevated-K+ epileptiform activity

Considerable variability in elevated-K+ epileptiform waveforms are observed across slices (Jensen and Yaari, 1988; Traynelis and Dingledine, 1988; Leschinger et al., 1993; Jensen and Yaari, 1997); we characterized elevated-K+ epileptiform activity in the CA1 region as having the following general characteristics after mechanical lesioning across the CA2 region. Two types of pre-ictal bursts were identified. “Fast-transition” pre-ictal activity started with a short train of low-amplitude high-frequency population spikes that immediately switched to high-amplitude high-frequency population spikes (Figure 1A, ER1); fast-transition pre-ictal activity propagated quickly (∼100 mm/s) across the CA1 region. “Slow-transition” pre-ictal like activity was characterized by a gradual increase in population spike and slow-field shift amplitude (Figure 1A, ER1; reminiscent of non-synaptic aggregate formation)(Bikson et al., 2003a); peak population spike size was generally smaller and activity more localized than that observed during fast-transition pre-ictal like activity.

**Figure 1:**
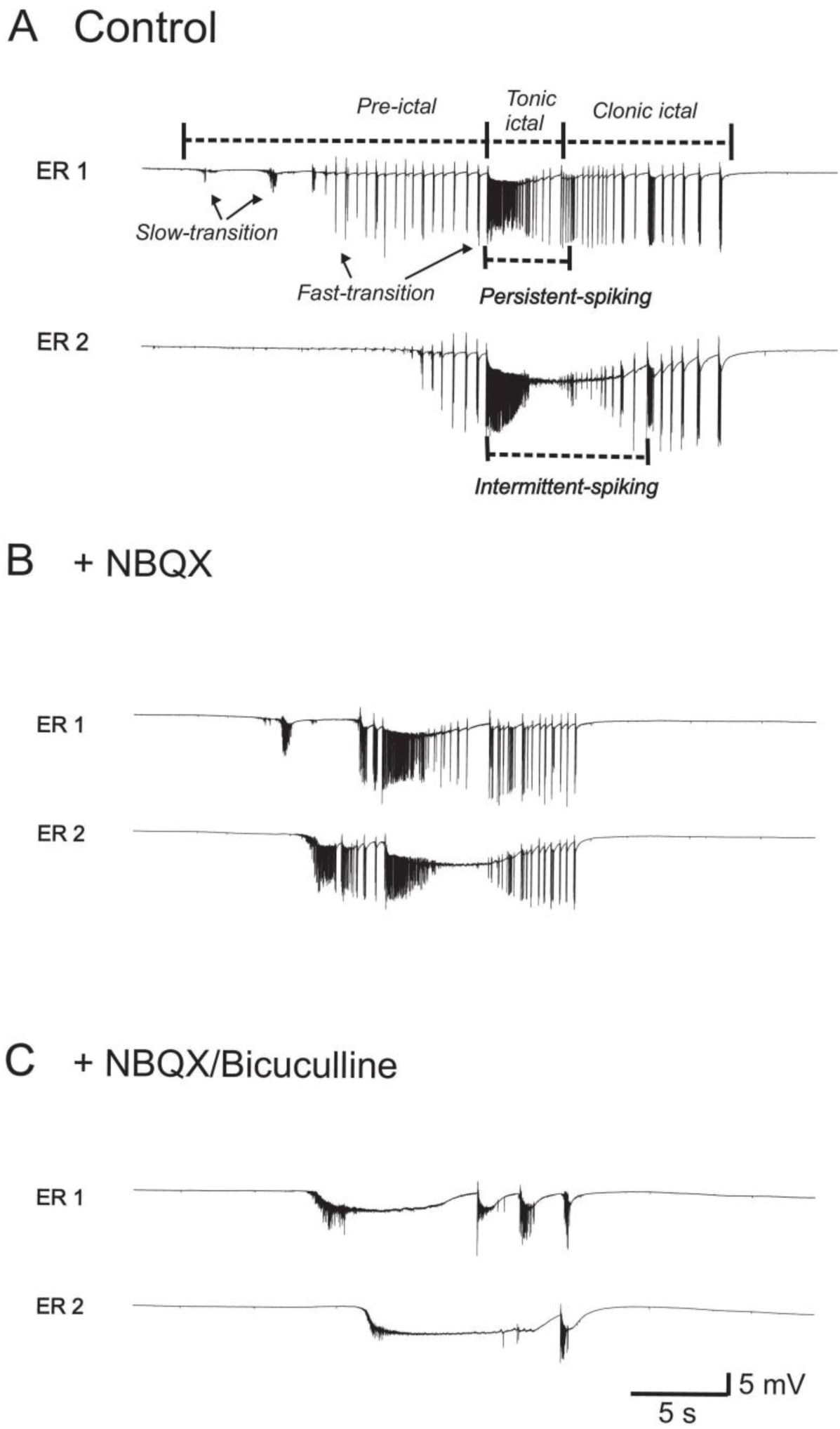
General waveform characterization of elevated-K+ epileptiform activity induced by potassium elevation in the CA1 region of hippocampal slices and effect of neurotransmitter receptor antagonists. (A) Extracellular field recordings with two electrodes (ER1, ER2) show distinct patterns of activity including pre-ictal activity preceding an ictal event. During the ictal phase population spikes could persist (“persistent-spiking”) or be transiently interrupted (“intermittent-spiking”). In this example, variation in waveform was observed across the CA1 region of the same slice. (B) Though altered, pre-ictal and both persistent- and intermittent-spiking ictal activity persisted in the presence of NBQX, an ionotropic glutamate receptor antagonist. (C) Further addition of ^bicuculline, a GABA^A^-receptor^ antagonist, blocked pre-ictal activity and changes ictal event waveform, notably suppressing population spike generation toward intermittent-spiking. These observations are consistent with the hypothesis that neither excitatory nor inhibitory function is required for ictal burst generation, but can shape ictal burst waveform (with reduced inhibition suppressing spiking) and the transition from pre-ictal to ictal activity

The period preceding the initiation of the ictal event was marked by an increase in the frequency and amplitude of pre-ictal events. Slow pre-ictal activity typically, but not necessarily, appeared first. In slices where fast-transition pre-ictal activity was observed, it was generally present immediately before ictal burst initiation and the initiation of ictal bursts often appeared to be triggered by a fast-transition pre-ictal event. Thus, in these cases, ictal events started with a rapid transition to large population spike activity and propagated quickly across the slice. In the remaining cases, where only slow-transition pre-ictal activity was observed or where no pre-ictal activity was present, ictal events generally started slowly (i.e. gradual increase in population spike amplitude) and propagated slowly across the slice (again reminiscent of non-synaptic seizure initiation, (Bikson et al., 2003a), though fast transition non-synaptic seizures are also observed (Haas and Jefferys, 1984)).

During the tonic phase of the ictal burst (defined as the duration of the slow negative field shift) repetitive large population spikes could persist for the entire duration of the ictal event or could be partially/completely disrupted for variable periods. As previously with non-synaptic activity, we classified tonic ictal events in which population spikes were disrupted (<1 mV) continuously for >2 s as “intermittent-spiking” ictal bursts (Figure 1A, ER2), and the remainder as “persistent-spiking” ictal bursts (Bikson et al., 2003a)

When the prolonged field shift returned to baseline, the “tonic” phase of the ictal event was defined as finished. In most slices, additional “clonic” discharges were observed (Traynelis and Dingledine, 1988). A quiescent period with no spontaneous field activity then followed.

Both the development and waveform of activity could vary significantly across the CA1 region though, as a general rule, large activity was more coherent across the slice. In pharmacological analysis, if different bursting types (i.e. intermittent and persistent spiking; Figure 1A) were observed at two locations in a slice then the effects of drugs on burst waveform at each site was considered separately.

### Role of glutamatergic and GABAergic synaptic function on elevated-K+ waveform

The role of excitatory neurotransmission in modulating elevated-K+ burst waveform was tested by adding ionotropic glutamate receptor antagonists. Addition of NBQX (20 µM; n=15), an AMPA/kainate-receptor antagonist, had no consistent effect on tonic ictal burst frequency (116 ± 49% control) or duration (111 ± 35% control). Addition of NBQX to slices showing persistent spike bursting (n=7; Figure 1B, ER1) did not induce a transition to intermittent spike bursting, increased peak population spike amplitude during the tonic ictal burst (128 ± 31% control; P<0.02). Addition of NBQX to slices showing intermittent spike bursting did not induce a transition to persistent-spiking in 8 of 9 cases (Figure 1B, ER 2); in the remaining slice a change to persistent-spiking bursts was observed following addition of NBQX. Subsequent addition of D-APV (25 µM; n=6), a NMDA-receptor antagonist, to slices still exhibiting intermittent-spiking, did not affect tonic ictal burst frequency (113 ± 10% control) or duration (98 ± 9 % control); it had no effect on intermittent spiking bursts in 4 of 6 slices (data not shown), but did induce a transition to persistent spiking in 2 slices. Fast transition pre-ictal like activity, slow-transition pre-ictal like activity, and clonic discharges (when present under control conditions) were not fully suppressed (blocked) by addition of NBQX and/or D-APV.

The role of inhibitory neurotransmission in modulating elevated-K+ burst waveform was tested by adding drugs that either block or enhance GABAergic function. Addition of ^bicuculline (^10^ µM), a GABA^A^-receptor antagonist, suppressed slow-transition pre-ictal^ activity in 7 slices, increased the frequency of tonic ictal discharges (161 ± 34 % control, p<.05), had no effect on tonic ictal burst duration (100 ± 37 % control) but ‘smoothed’ the tonic burst negative field shift (decreased population spikes during ictal bursts) - one slice generated spreading depression and was excluded. In those slices showing persistent spiking bursts under control condition (n=6), bicuculline induced transition to intermittent-spiking ictal bursting (data not shown). A similar effect was observed when bicuculline (10 µM) was added with NBQX (Figure 1C) and D-APV (data not shown) in 8 of 9 slices (Figure 1), in the remaining slice, the discharge type remained persistent-spiking.

The addition of CGP ^55845^ (^2^.^5^-^150^ µM, n=^10^), a GABAB^-receptor^ antagonist (Davies et al., 1993), had little effect on ictal or preictal discharges. There was no consistent effect on burst duration (116 ± 22% control); burst frequency was unchanged (99 % control); persistent-spiking bursting became intermittent in only one out of three slices and intermittent-spiking bursts remained intermittent (n = 7)(data not shown).

Addition of nipecotic acid (0.5-1 mM, n=9), a GABA uptake blocker (Krogsgaard-Larsen and Johnston, 1975), to slices showing intermittent spiking bursting, had no effect on tonic ictal burst duration (104 ± 25% control), no significant change in tonic ictal burst frequency (79 ± 11% control), caused a transition to persistent-spiking bursts, and promoted the generation of pre-ictal activity (Figure 2A) in 6 of 9 slices; the remaining 3 slices were excluded because in 2 ictal activity stopped, leaving only inter-ictal activity and in one slice spreading depression was induced. Further perfusion with higher concentration of nipecotic acid (1.5-3 mM, n=6) either 1) induced a transition back to intermittent-spiking bursting and suppressed clonic bursts; and/or 2) induced spreading depression (Figure 2). Addition of a different GABA uptake blocker, NNC-711(Suzdak et al., 1992) (2.5-50 µM), supported the results obtained with nipecotic acid. Slices exhibiting intermittent-spiking bursts showed a transition to persistent-spiking activity (n=2; Figure 2B), a transition to only pre-ictal activity (n=3), or the generation of spreading depression (n=5).

**Figure 2:**
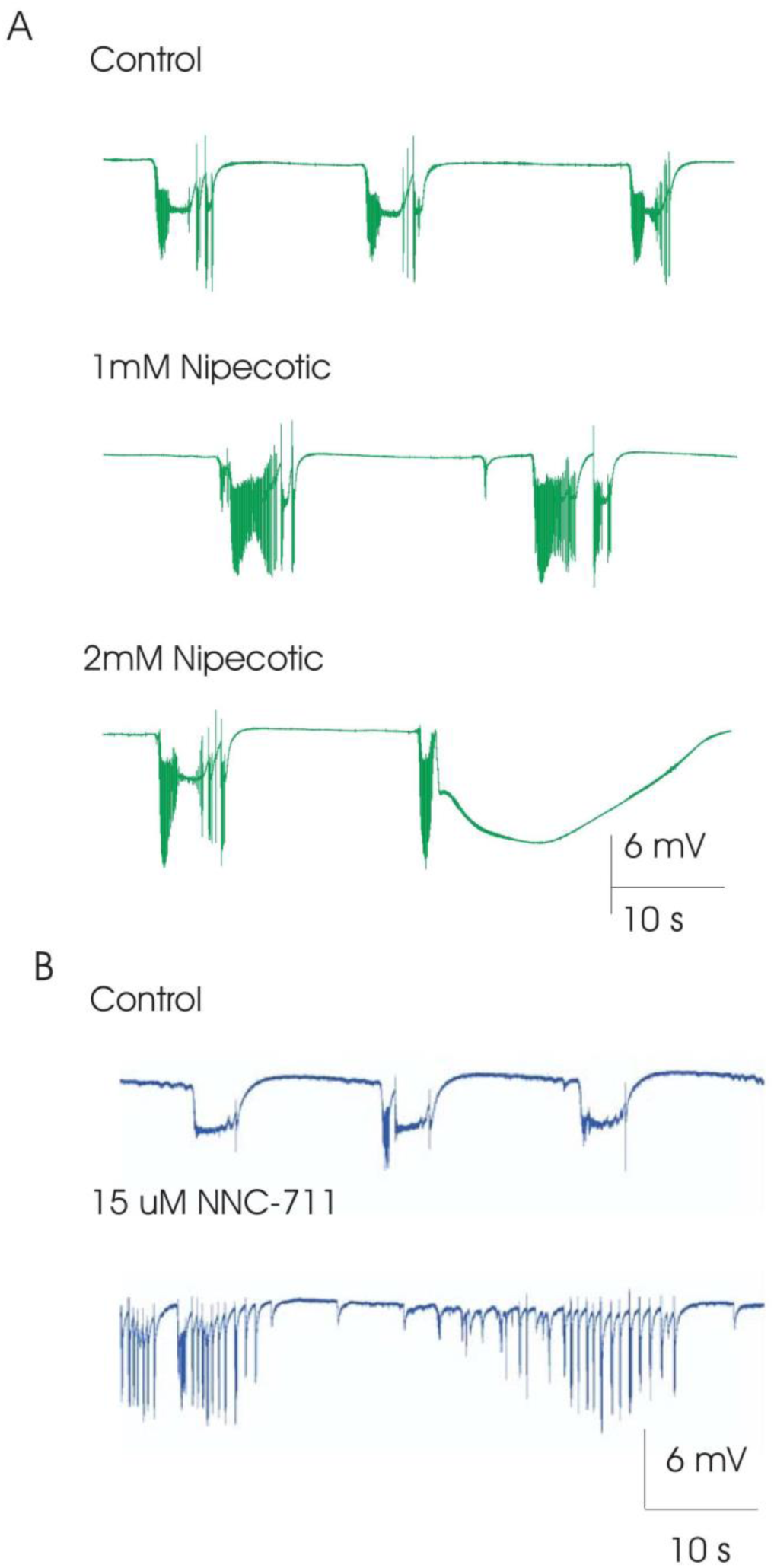
Effect of GABA-uptake inhibitors on elevated-K+ induced epileptiform activity waveform. (A) Addition of 1 mM nipecotic acid, a GABA-uptake blocker, promoted the generation of population spikes during ictal activity, converting intermittent-spiking ictal activity to persistent spiking while promoting pre-ictal activity. Increasing nipecotic acid dose from 1 to 2 mM resulted in spreading depression like activity. (B) Addition of 15 uM NNC-771, also a GABA-update blocker, could similarly convert intermittent-spiking ictal activity to persistent spiking, while enhancing pre-ictal activity. These observations are consistent with the hypothesis that enhancing GABA function (enhancing inhibition), promotes neuronal spiking during ictal busts while inhibiting the transition from pre-ictal to ictal activity.

To test the potential role of depolarizing GABA in the transition to epileptic seizures we perfused the slice with ethoxyzolamide (100 µM ethoxyzolamide dissolved in 0.1 M NaOH), which has been shown to block the depolarizing effect of GABA. In 15 of 16 slices, ethoxyzolamide produced gradual decrease seizure frequency followed by cessation of seizure-like activity (Figure 3). In 1 of 16 slices, a transition from persistent to intermittent ictal activity was produced with a block of the clonic phase. Washout of ethoxyzolamide was associated with reoccurrence of seizure activity. In 4 slices where pre-ictal activity was present, preictal activity persisted after wash in of ethoxyzolamide while ictal activity was blocked. Three slices were perfused only with 0.1 M NaOH to exclude the possibility that the above mentioned observation was due to its effect. Perfusion of slices with 0.1 M NaOH had no effect of epileptic activity.

**Figure 3:**
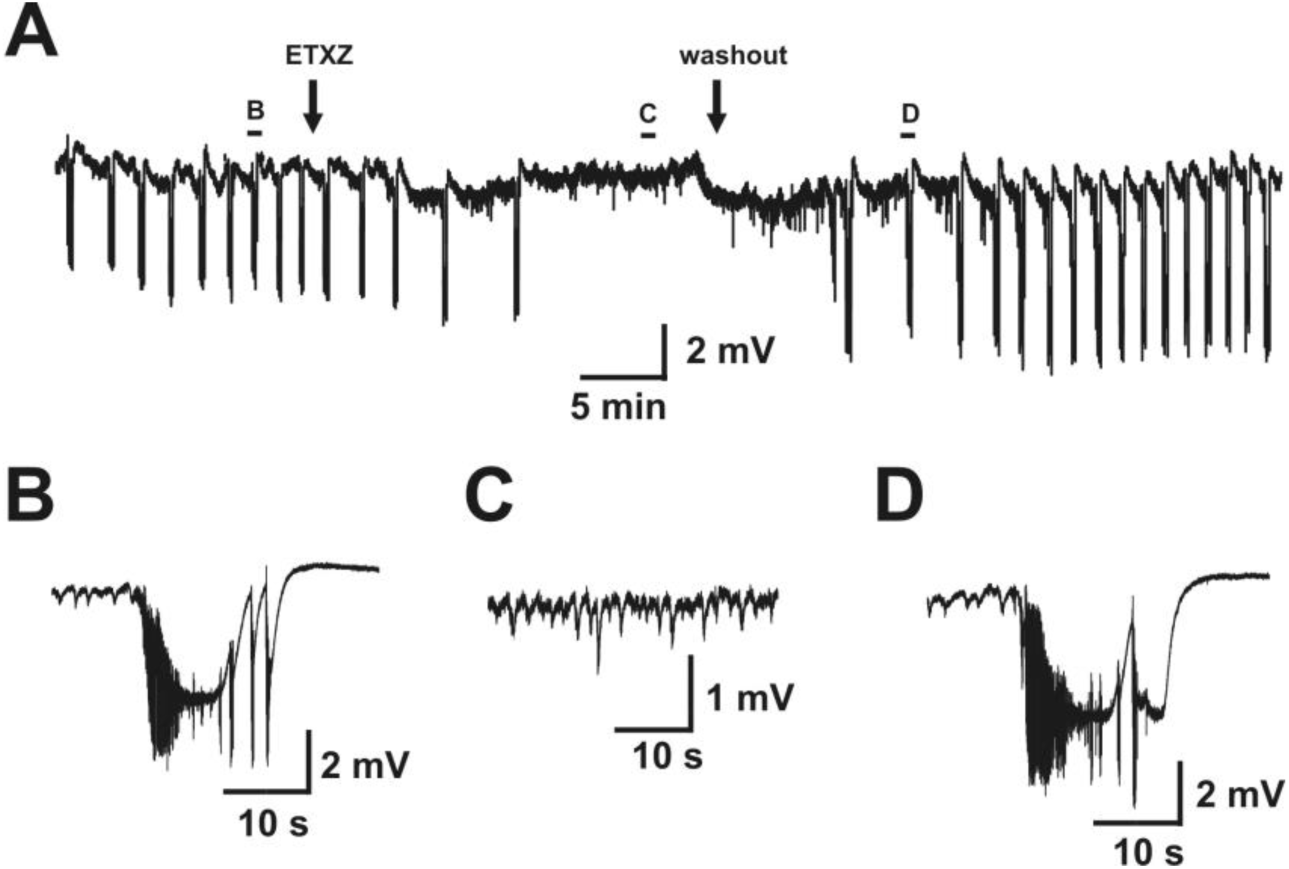
Block of depolarizing GABA by ethoxyzolamide (ETXZ). (A) Recording from CA1 shows spontaneous repeated seizures characterized by repeated population discharges, which are superimposed on a large direct current (DC) shift (B). ETXZ causes decreased seizurefrequencyand disappearance of seizure activity (C). Washout of ethoxyzolamide leads to reoccurrence of epileptic seizures.

In summary, multifaceted changes in ictal and interictal activity were brought about ^by modifying GABAergic synaptic function. The results are consistent with a role forGABA^A release inpromoting the generation of pre-ictal activity (which may be preventing the transition to ictal activity), reducing ictal burst generation frequency, and modulating the ^generation of population spikes during ictal bursts. Antagonizing GABA^B ^function had^ minimal effect, moderately modulating burst duration. While the nonspecific effects of ethoxyzolamide limit definitive conclusions on the role of depolarizing GABA, suppression of ictal but not pre-ictal activity by ethoxyzolamide along with the distinct effects of low and high concentrations of nipecotic acid (a GABA uptake blocker) support a multi-dimensional role for GABAergic synaptic function. Despite a modulatory role, inhibitory synaptic function is not necessarily critical for ictal activity. By contrast, glutamate receptor antagonists had no consistent effect on pre-ictal activity, tonic ictal burst frequency, or clonic burst activity, though in some slices antagonizing glutamate receptors enhanced population spike generation during ictal activity.

### Intracellular activity during elevated-K+ bursting

The source of population spike disruption during intermittent-spiking bursts was investigated using sharp intracellular recording of CA1 pyramidal neurons. A total of 7 cells (from 7 rats) were recorded during intermittent spiking (4 under control conditions, 3 in the presence of bicuculline). All neurons recorded depolarized for the entire duration of the ictal event. Interruptions in population spike generation were associated with stoppages in individual cell firing (Figure 4).

**Figure 4:**
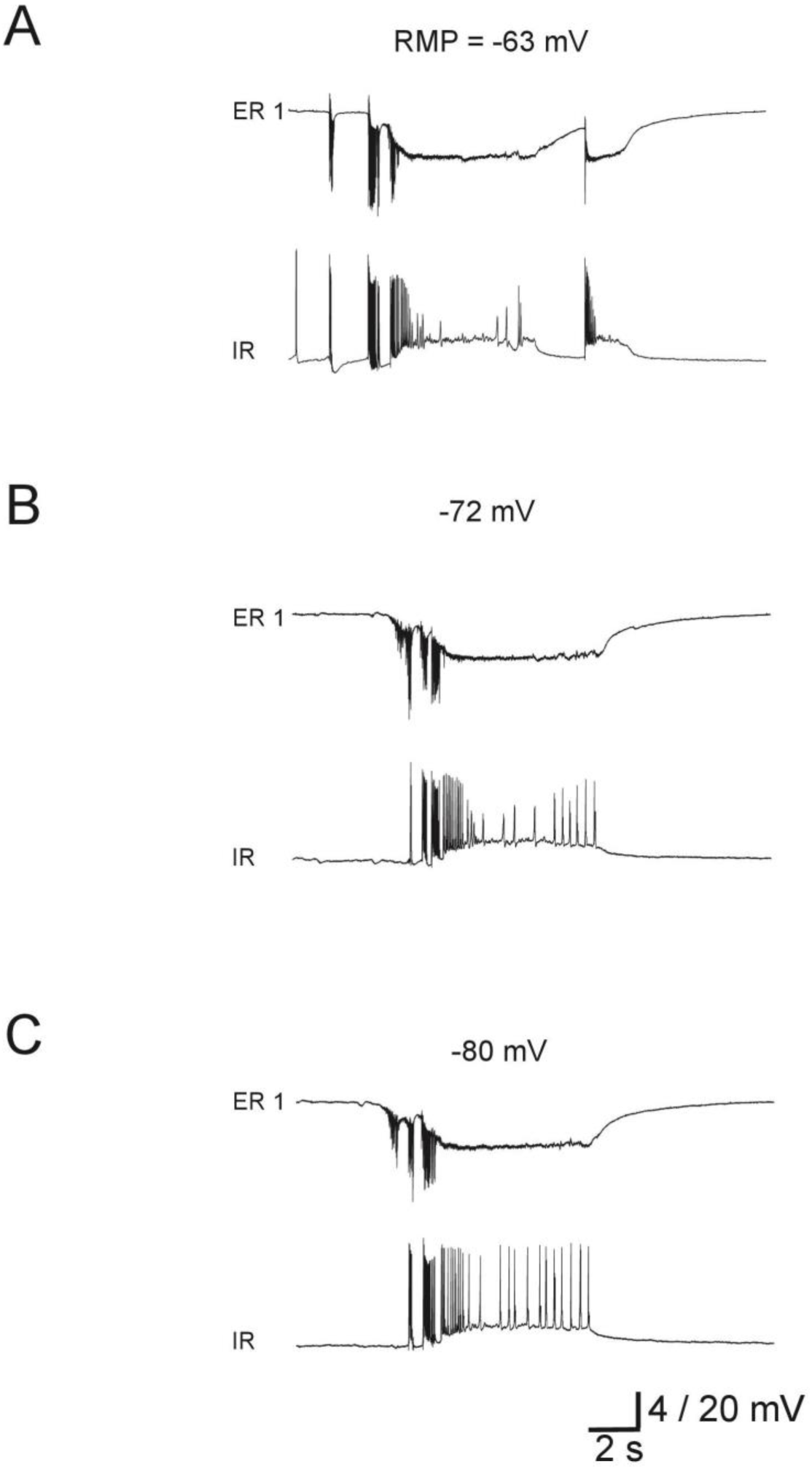
Concurrent extracellular field recording (ER) and intracellular recording (IR) from a CA1 neuron during elevated-K+ epileptiform activity. Suppression of population spikes in the field recording (intermittent spiking) was associated with a depolarization of the neuron and suppression of action potential generation. Intracellular injection of hyperpolarizing (-0.2 nA) current, but not depolarizing (0.2 nA) current, promoted the generation of action potentials by the neuron, even while population spikes remained suppressed. Repeated steps of 0.5 nA were concurrently used to measure cell resistance. These results support the hypothesis that interruption of population spikes during an ictal event (intermittent spiking) reflect depolarization block of neurons.

To investigate if depolarization block accounted for this stoppage, neurons were hyperpolarized by steady current injection (n=6; 4 under control conditions, 2 in the presence of bicuculline). In all neurons tested, hyperpolarization enhanced ‘excitability’ during the burst as indicated by the generation of spontaneous action potentials during the entire ictal burst or by a reduced threshold to action potential initiation by a brief intracellular depolarizing step (Figure 4); these responses to hyperpolarization are consistent with an interruption in firing during intermittent spiking burst reflecting depolarization block of action potential initiation.

Changes in membrane resistance during bursts were monitored with brief hyperpolarizing pulses (n=6 cells). During intermittent-spiking bursts membrane resistance was 29 ± 9 M Ω pre-burst and 22 ± 10M Ω during a burst (average change 6.6 ± 6.5 M Ω and 75 ± 25% burst/inter-burst; P=0.056). These values are comparable with previous reports (Traynelis and Dingledine, 1988; Jensen and Yaari, 1997). We found that hyperpolarizing neurons from rest to -80 mV by injection of a constant current had no significant effect on membrane resistance between bursts (average 25 ± 8 M Ω); with this holding current membrane resistance trended toward an increase during intermittent-spiking bursts to 29 ± 10 M Ω (average delta 4.1 ± 3.4 M Ω, 117 ±13% burst/inter-burst; P=0.056). The membrane resistance during a burst when constant current was injected was 143 ± 33% that of the resistance during a burst without constant current injection.

## Discussion

Synaptic mechanisms underlie all brain activity including pathological activity. Yet non-synaptic mechanisms have been shown to be sufficient to generate electrographic seizures after synaptic activity is blocked. We addressed how in the presence of synaptic activity, both synaptic and non-synaptic mechanisms contribute to seizure initiation, and dynamics. We categorized this dependence through selective blockade and promotion of synaptic mechanisms during elevated-K+ induced electrographic seizures in CA1 of hippocampal slices. We confirm that ictal activity can persist after blocking excitatory and/or inhibitory synaptic transmission. Block of excitatory synaptic transmission had minimal effects on ictal burst waveform. Modulation of inhibitory synaptic transmission had a multifaceted influence on both pre-ictal and ictal activity, as well as population spikes during the ictal event. In some cases, inhibitory synaptic transmission enhanced aspects of epileptiform activity. We outline a framework for electrographic seizure genesis in CA1 based on successful or failed transition from pre-ictal events, synaptic transmission mediates this transition and influences seizure dynamics (Figure 5).

**Figure 5:**
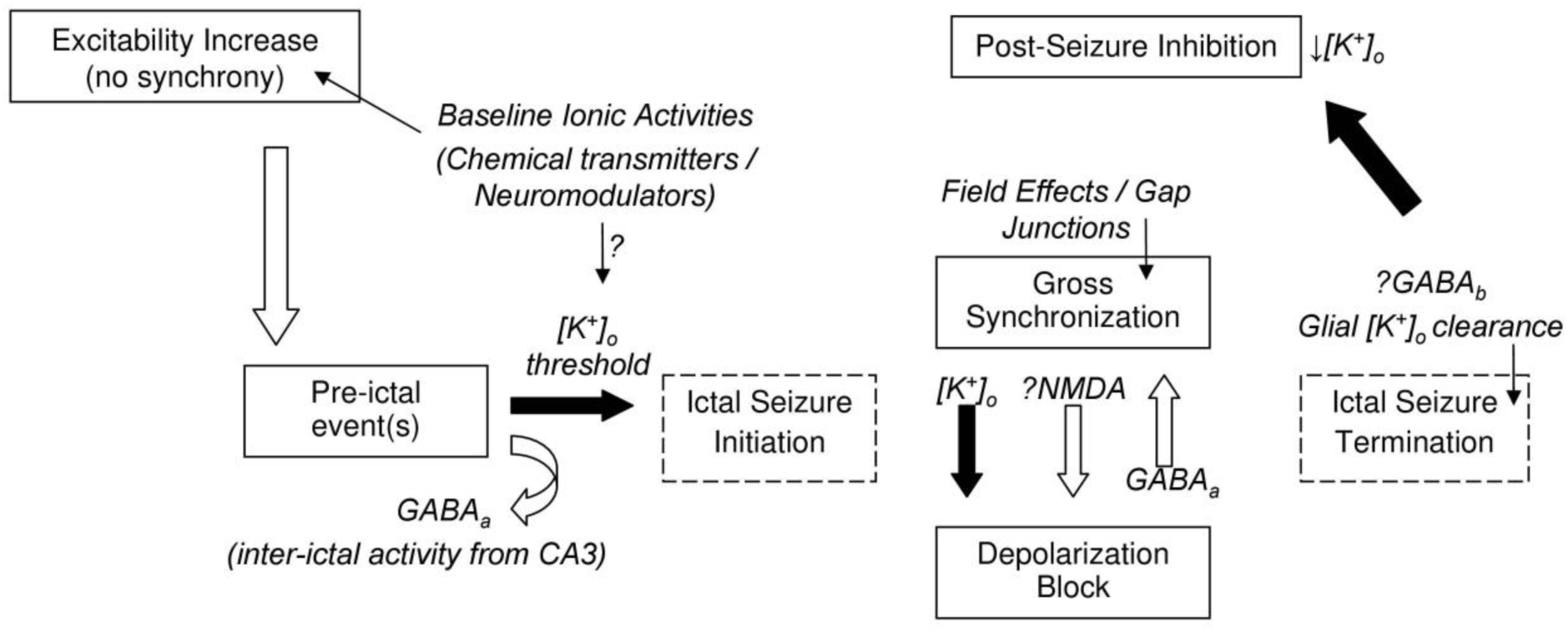
Schematic of proposed general frame-work for icto-genesis in the CA1 region of the hippocampus. Changes in baseline ionic or neuro-modulator activities (either exogenous/applied or transient endogenous) ^result in an initial increase in excitability. Population activity is first observed as a pre-ictal event; GABA^A function may prevent an immediate transition to ictal bursting. Non-synaptic mechanisms are sufficient to ^initiate, maintain, and terminate a seizure, with [K+]^o ^dynamics presumably plating a key role. Synaptic^ mechanisms may affect population spike size by moving neurons in/out of depolarization block but may not directly contribute to fast synchronization. The precise mechanisms leading to seizure termination, when significant synchronization is still observed, remain unclear. The post-seizure inhibition is followed by a gradual increase in excitability. According to this scheme, the primary role of chemical neuro-modulators is in regulating excitability levels and effecting epileptiform activity initiation rate, and in shaping ictal event waveform.

## Electrographic seizure initiation from pre-ictal events

We propose electrographic seizures arise from locally generated pre-ictal activity (Figure 1A). Previous reports showed that CA1 can both locally generate (what we call) preictal activity (Tancredi and Avoli, 1987), and propagate inter-ictal synaptic drive from CA3 (Jensen and Yaari, 1988; Traynelis and Dingledine, 1988). CA3 inter-ictal activity is remote and upstream from CA1, and persists during the ictal events in CA1. CA3 inter-ictal activity may inhibit icto-genesis in CA1. We propose pre-ictal activity in CA1 is not simply correlated with icto-genesis (Cymerblit-Sabba and Schiller, 2010; Jiruska et al., 2010; Zhang et al., 2012; Fujita et al., 2014; Perucca et al., 2014) but reflects the successful or aborted initiation of an ictal event. Synaptic function influences this transition. Notably, modulation of GABA_A_ ynaptic function affected the occurrence of pre-ictal activity and seizure frequency inversely (GABA_A_ receptor antagonist bicuculline increased ictal discharge frequency and abolished pre-ictal activity, while GABA uptake blocker nipecotic acid decreased ictal frequency and promoted pre-ictal activity) consistent with the hypothesis that GABA_A_ inhibits the transition from pre-ictal to ictal activity. Both crossing of a seizure threshold during a pre-ictal event and ictal event dynamics may be driven by non-synaptic mechanisms such as extracellular K^+^ ([K^+^]_o_) accumulation (Nelken and Yaari, 1987; Jensen and Yaari, 1997; Kager et al., 2000).

In our model, glutamatergic synaptic function does not play a critical role in facilitating pre-ictal to ictal transition, though this may not generalize to other conditions or brain regions (Patrylo et al., 1994; Bragin et al., 2009; Huberfeld et al., 2011). However, our observations with GABA modulation caution that perturbations that appear to inhibit pre-ictal activity may in fact accelerate transition to electrographic seizures. This should be considered across pharmacological studies of epilepsy.

## Synchronization and depolarization block during electrographic seizures

Across clinical cases and animal models, the development of pre/inter-ictal activity is reported to correlate with seizure dynamics (Huberfeld et al., 2011; Zhang et al., 2012; Perucca et al., 2014). Our results are consistent with the hypothesis that the spatio-temporal synchronization of neurons at the start of seizures depends on the preceding pre-ictal activity; with fast- or slow-transition pre-ictal activity leading to similar electrographic seizure synchronization/propagation patterns. These can be driven by slow (aggregate formation (Bikson et al., 2003a) and K^+^ diffusion (Lian et al., 2001; Durand et al., 2010; Martinet et al., 2017)), and fast (field effects (Haas and Jefferys, 1984; Qiu et al., 2015)) non-synaptic processes. Our results do not suggest a strong influence of synaptic function on synchronization except through mediation of depolarization block.

Depolarization block is prevalent in a wide class of epilepsy models ((Bikson et al., 2003b); see also “start-stop-start” and “electrodecremental” events; (Blume and Kaibara, 1993)). Depolarization block was commonly observed during elevated-K^+^ ictal discharges, influenced by [K^+^]_o_ transients which peak in the tonic ictal phase (Jensen and Yaari, 1997; Bikson et al., 2002). In the elevated-K^+^ model, as in other electrographic seizures (Bikson et al., 2003b) the absence of neuronal firing does not herald the termination of the ictal discharge nor does it necessarily affect their duration. Modulation of either inhibitory and excitatory synaptic function had no consistent effect on tonic ictal burst duration. The termination of non-synaptic ictal discharges may depend on K^+^ clearance mechanisms (Bikson et al., 1999). A decrease in [K^+^]_o_ coincides with the termination of the tonic phase of elevated-K^+^ discharges (Traynelis and Dingledine, 1988; Jensen and Yaari, 1997); a moderate K^+^ rise is maintained for the duration of the clonic phase.

The initially counter-intuitive enhancement of population spike size (synchronization) after antagonism of glutamatergic-synaptic function and suppression of population spikes after antagonism of GABAergic function is consistent with influencing depolarization block. Though our results support a role for hyperpolarizing GABA in preventing depolarization block during control elevated-K^+^ bursting, our finding that increasing concentration of nipecotic acid could revert bursting back to intermittent-spiking may indicate that, under certain conditions, GABA can exert a depolarizing influence (Bracci et al., 1999). Our findings with ethoxyzolamide may suggest a role for depolarizing GABA in transition from ^pre-ictal to ictal activity. Finally, synaptic activity and [K^+^]^o ^are intricately linked including^ through co-transporter activity (Malenka et al., 1981; Hubner et al., 2001; Bihi et al., 2005; Kaila et al., 2014). Nonetheless, even in the presence of synaptic function, depolarization block would indicate non-synaptic governing of seizure dynamics, since [K^+^]_o_ accumulation would now play a central role in seizure maintenance.

## A single theoretical framework for ictogenesis in the hippocampal formation?

Various models of hippocampal electrographic seizure generation differ in their reported response to synaptic antagonists. One possibility is that these models represent fundamentally different mechanisms of seizure development. Alternatively, all these models can be explained by a single nuanced framework where non-synaptic mechanisms underpin seizure development and synaptic mechanisms modulate seizure initiation and waveform.

The frequency of ictal burst depends on neuronal excitability (reviewed in (Bikson et al., 1999)). By simply influencing “general” excitability, synaptic function can alter ictal burst frequency - which when changed above/to zero results in qualitative seizure control. Consistent with this proposition, we found that modulation of GABA function could affect ^ictal burst frequency and, in separate experiments, that addition of a GABA^A ^receptor^ antagonist to slices showing only pre/inter-ictal activity could induce the generation of ictal bursts (not shown). In the dentate gyrus, enhancement on neuronal excitability by *either* ^elevating [K^+^]^o^, decreasing [Ca^2^+]^o^, or blockade of GABA^A ^receptors is necessary to generate^ ictal bursts (Patrylo et al., 1994). In CA1, elevated-K^+^ ictal bursting becomes dependent on synaptic function when [Ca^2^+]_o_ and extracellular Mg^2^+ concentrations ([Mg^2^+]_o_) are increased (Traynelis and Dingledine, 1989); divalent cations reduce neuronal excitability by ‘charge screening‘. Low-Ca^2^+ bursting is suppressed by exogenous GABA only after [Ca^2^+]_o_ is moderately raised (Watson and Andrew, 1995). Across models, any factors that reduce neuronal excitability, including temperature (Leschinger et al., 1993; Schuchmann et al., 2002), animal age (Gloveli et al., 1995), slice size/thickness (Traynelis and Dingledine, 1989), and superfusate level (Schuchmann et al., 2002) could thus *induce* sensitivity to neurotransmission.

Support for a role for neurotransmitters in modulating the rate of ictal burst initiation (frequency) also comes from studies on non-synaptic ictal bursting (where endogenous neurotransmitter release is blocked) in which exogenous application of neuromodulators and neurotransmitters that increase (DL-homocysteic acid, histamine) or decrease (GABA, taurine, adenosine) excitability preferentially modulated burst frequency (Haas and Jefferys, 1984; Lee et al., 1984; Watson and Andrew, 1995; Xiong and Stringer, 2001). Application of ^either exogenous GABA or baclofen (a GABA^B ^receptor agonist) also promotes the^ generation of ‘pre-ictal’ activity and *increases* population spike amplitude during non-synaptic bursting (Watson and Andrew, 1995; Xiong and Stringer, 2001)

The dependence of ictal burst frequency on neuronal excitability (Fox et al., 2007) could reflect the need for a critical mass of rapidly firing neurons to induce the regenerative potassium build-up correlated with all ictal bursts (Jefferys, 1995). In the 4-AP model, increasing [Ca^2^+]_o_ and/or [Mg^2^+]_o_, which reduces excitability by increasing charge screening, induces a dependence on ionotropic synaptic function (Voskuyl and Albus, 1985; Martin et al., 2001; Barbarosie et al., 2002); this dependence, however, relates in-turn to non-synaptic potassium accumulation (Barbarosie et al., 2002). In the low-chloride model, dependence on synaptic function may be overcome by increasing neuronal excitability either by increasing stimulation intensity (Avoli et al., 1990) or by increasing [K^+^]_o_ (Demir et al., 1999). Clearly, in models which inherently depend on enhancement of excitatory neurotransmission; antagonism of the targeted synaptic pathway (though not necessarily other synaptic pathways) would reverse the enhancement of neuronal excitability (Anderson et al., 1986; Neuman et al., 1988).

Inter-ictal activity is generally dependent on synaptic mechanisms and can both promote and suppress ictal bursting (Khosravani et al., 2003); this will, therefore, indirectly complicate the pharmacological profile of ictal bursting (Swartzwelder et al., 1987; Bragdon et al., 1992). It is also necessary to consider the interaction of synaptic mechanisms with non-synaptic function (in particular K^+^ regulation mechanisms) in interpreting the effects of neurotransmitter modulators on ictal bursts (Pumain et al., 1985; Martin et al., 2001; Thuault et al., 2002)

In summary, we propose that if the development of ictal activity in the hippocampus is dependent on non-synaptic regenerative K^+^ accumulation, seeded by pre-ictal events (Fertziger and Ranck, 1970; Nelken and Yaari, 1987), it follows that neurotransmission ^would act primarily to influence the chance (the frequency) a threshold [K^+^]^o ^is reached. Synaptic function, along with [Ca^2^+]^o^, may also influence the [K^+^]^o ^threshold value. One^ initiated, seizures can then be modulated by, but do not require, neurotransmission. The combination of increased excitability and reduced synaptic inhibition is not sufficient to generate ictal discharges (Tancredi and Avoli, 1987; Bikson et al., 2002) as feed-forward mechanisms for K^+^ accumulation are necessary.

## Acknowledgements

This worked was funded in part by MRC grant (#G9901454) Epilepsy Research UK grant to JGRJ and JF, the Wellcome Trust Programme Grant to JGRJ, and DoD AFOSR grant (#FA9550-13-1-0073) and NIH/NSF grant to MB (#5R01MH092926).

